# Suppression of aminoglycoside-induced premature termination codon readthrough by the TRP channel inhibitor AC1903

**DOI:** 10.1101/2021.04.07.438788

**Authors:** Alireza Baradaran-Heravi, Claudia C. Bauer, Isabelle B. Pickles, Sara Hosseini-Farahabadi, Aruna D. Balgi, Kunho Choi, Deborah M. Linley, David J. Beech, Michel Roberge, Robin S. Bon

## Abstract

Nonsense mutations, which occur in ~11% of patients with genetic disorders, introduce premature termination codons (PTCs) that lead to truncated proteins and promote nonsense-mediated mRNA decay. Aminoglycosides such as gentamicin and G418 permit PTC readthrough and so may address this problem. However, their effects are variable between patients, making clinical use of aminoglycosides challenging. In this study, we addressed the hypothesis that TRP non-selective cation channels contribute to the variable effect of aminoglycosides by controlling their cellular uptake. To attempt to identify the channel type involved, we tested AC1903, a 2-aminobenzimidazole derivative recently reported to selectively inhibit TRPC5 cation channels. AC1903 consistently suppressed G418 uptake and G418-induced PTC readthrough in the DMS-114 cell line and patient-derived JEB01 keratinocytes. In an effort to validate the suggested role of TRPC5, we tested an independent and more potent inhibitor called Pico145, which affects channels containing TRPC1, TRPC4 and TRPC5 but not other TRPCs or other channels. Unexpectedly, Pico145 was completely without effect, suggesting that AC1903 may work through other or additional targets. Consistent with this suggestion, AC1903 inhibited multiple TRPC channels including homomeric TRPC3, TRPC4, TRPC5, TRPC6 as well as concatemeric TRPC4–C1 and TRPC5–C1 channels, all with low micromolar IC_50_ values. It also inhibited TRPV4 channels but had weak or no effects on TRPV1 and no effect on another non-selective cation channel, PIEZO1. Overall, our study reveals a suppressor of aminoglycoside-mediated PTC readthrough (i.e., AC1903) but suggests that this compound has previously unrecognised effects. These effects require further investigation to determine the molecular mechanism by which AC1903 suppresses aminoglycoside uptake and PTC readthrough.

## Introduction

Nonsense mutations account for ~11% of all genetic lesions in patients with inherited diseases (1). They introduce a TAG, TGA or TAA premature termination codon (PTC) and typically result in production of mRNAs with decreased stability as well as defective truncated proteins. PTC readthrough is a mechanism by which ribosomes recognise nonsense mutations as sense codons, enabling synthesis of the full-length and functional protein rather than truncated product. Aminoglycoside antibiotics were the first and are still among the most active chemicals discovered to induce PTC readthrough in yeast (2), mammalian cells (3), animal models (4, 5), and patients (6, 7). These compounds bind at the decoding centre of eukaryotic ribosomes and facilitate pairing of near-cognate aminoacyl-tRNAs to the PTCs resulting in formation of full-length protein (8, 9). Several non-aminoglycoside readthrough compounds have also been identified, including negamycin, tylosin, RTC13, RTC14, GJ71, GJ72 and ataluren (10–14). However, these compounds induce PTC readthrough at low rates and often at or below the detection limit of western blotting for endogenous protein expression.

In addition to their severe in vivo toxicity, aminoglycosides induce variable levels of PTC readthrough in cell lines as well as patients, making long-term clinical administration of these drugs challenging (15–20). In general, variation in drug response is caused by differential local drug concentrations (pharmacokinetics) or drug actions (pharmacodynamics) and to a great extent is attributed to genetic variations (21). Variations in aminoglycoside-induced PTC readthrough could be in part related to the PTC and its surrounding sequence as well as its distance to the poly-A tail sequence. Moreover, differential cellular uptake of aminoglycosides and consequent variable intracellular concentrations of these compounds may underlie mechanisms of PTC readthrough variation. However, the correlation of genetic variations in genes involved in cellular uptake of aminoglycosides and their contribution to PTC readthrough in human cells is not well understood.

Endocytosis (22, 23) and permeation through non-selective cation channels (including TRP channels) (24–28) are the main proposed mechanisms for cellular uptake of aminoglycosides. In this study, we addressed the hypothesis that TRP channels may contribute to variable effects of aminoglycoside-mediated PTC readthrough. Following the observation that low PTC readthrough in response to treatment with the aminoglycoside G418 (which is cationic at physiological pH) was correlated with a mutation in *TRPC5*, we tested the effects of the 2-aminobenzimidazole derivative AC1903, recently reported as a selective TRPC5 channel inhibitor. We found that AC1903, but not the selective TRPC1/4/5 inhibitor Pico145, suppresses cellular uptake of the aminoglycoside G418 and G418-induced PTC readthrough. Consistent with these results, we subsequently found that AC1903 is not a selective inhibitor of TRPC5 (or TRPC1/4/5) channels, but inhibits multiple TRP channels, including TRPC3, TRPC4, TRPC6 and TRPV4. Our work reveals AC1903 as a suppressor of aminoglycoside-induced PTC readthrough, but highlights that the mode-of-action is not understood, and may enable new investigations into the use of aminoglycosides for the treatment of genetic diseases. In addition, this study underlines that investigations of TRPC5 channel biology with non-selective inhibitors such as AC1903 need to be supported carefully by orthogonal chemical or genetic approaches.

## Results

### Variable aminoglycoside-induced PTC readthrough in cell lines with identical TP53 nonsense mutation

Variation in PTC readthrough response has been reported among individuals or cell lines with different nonsense mutations. Although this in part highlights the importance of the stop codon and its surrounding sequence as well as the physical location of the nonsense mutation in the mRNA sequence, understanding the contribution of other underlying genetic variations becomes more complicated. To rule out the effect of the sequence and position of nonsense mutations on response variation to potential PTC readthrough modulators, we studied two cancer cell lines, DMS-114 and TC-71, with identical homozygous nonsense mutation R213X in their *TP53* gene. Exposure of DMS-114 cells to increasing concentrations of the aminoglycoside G418 for 24 h resulted in strong, dose-dependent increase in production of full-length p53 (the readthrough product) up to 174-fold compared to untreated cells (**Figure 1A**). In contrast, increasing concentrations of G418 only elicited a small increase in PTC readthrough in TC-71 cells, up to 9-fold compared to untreated cells (**Figure 1A**). To find out whether the reduced response to G418 in TC-71 cells was associated with lower intracellular levels of G418, we measured concentrations of G418 in both cell lines at 0, 8 and 24 h post exposure to G418 (100 μg/ml). Compared to DMS-114, intracellular G418 levels were slightly but significantly lower in TC-71 cells at 8 h and 24 h post G418 exposure, by ~12% and ~4% respectively (**Figure 1B**). These data suggest that reduced intracellular G418 level in TC-71 cells vs. DMS-114 may be partially responsible for the difference in G418-induced PTC readthrough.

**Figure 1.**
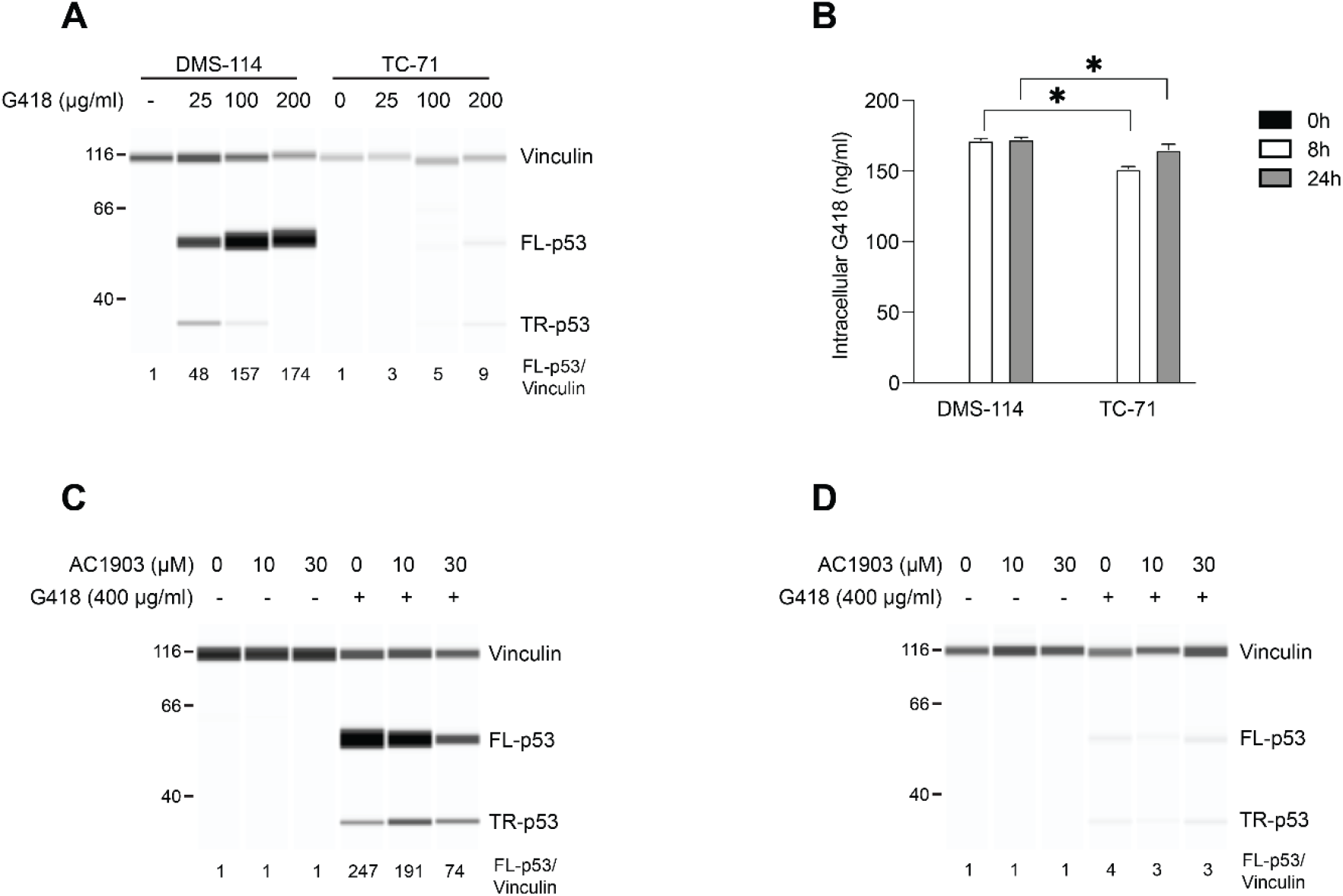
Variable PTC readthrough in cell lines with the same homozygous *TP53* nonsense mutation. A) DMS-114 and TC-71 cell lines with homozygous *TP53* nonsense mutation (R213X) were exposed to indicated concentrations of G418 for 24 h and p53 levels (full-length, FL-p53; truncated, TR-p53) were measured. B) 100 μg/ml G4180 was added to DMS-114 and TC-71 cells and its total intracellular levels were measured at indicated time points in duplicates. C,D) DMS-114 (C) or TC-71 (D) cells were pretreated with indicated concentrations of AC1903 for 3 h followed by exposure to 400 μg/ml G418. After 3 h medium was replenished and at 24 h p53 levels were measured. In panels (A, C, D) protein levels were measured using automated capillary electrophoresis western analysis and vinculin was used as loading control. * indicates statistically significant difference between samples (P < 0.05).

To find out the possible contribution of genetic variants to this variation we searched exome sequencing data in COSMIC (https://cancer.sanger.ac.uk/cosmic) and identified a hemizygous p.R175H missense mutation in *TRPC5* in the TC-71 cell line not present in the DMS-114 cell line. This gene encodes the protein TRPC5, which forms homo- and heterotetrameric, non-selective cation channels permeable by Ca^2+^ and Na^+^ (29–34). Based on several pathogenicity prediction algorithms including SIFT (35) and PolyPhen-2 (36), this mutation is considered tolerable and benign. R175 of TRPC5 is located in a putative zinc-binding domain – consisting of H172, C176, C178 and C181 – that is conserved between TRPC proteins (37, 38). This domain has also been implicated in TRPC5 regulation by S-glutathionylation (39), as well as S-palmitoylation required for TRPC5 trafficking to the plasma membrane (40). Because other non-selective cation channels of the TRP family have been reported to contribute to aminoglycoside import (24–28), we decided to test the potential role of TRPC5 in this process.

### AC1903 suppresses G418 trafficking and PTC readthrough

To investigate the potential role of TRPC5 in modulation of G418 intracellular levels and G418-induced PTC readthrough, we exposed DMS-114 and TC-71 cells to various concentrations of AC1903, a recently reported TRPC5 inhibitor (41, 42). Cells pre-treated with AC1903 for 3 h were exposed to 400 μg/ml G418 for another 3 h, followed by media replacement and incubation for another 18 h. Exposure to AC1903 alone did not affect PTC readthrough in either cell line (**Figure 1C–D**). As expected, G418 elicited strong PTC readthrough in DMS-114 but not in TC-71 cells (**Figure 1C–D**). Interestingly, in DMS-114 but not in TC-71 cells, combination of AC1903 (10 or 30 μM) with G418 reduced PTC readthrough by 23% and 70%, respectively, compared to G418 alone (**Figure 1C–D**).

In order to better understand the effects of AC1903 on G418 uptake and PTC readthrough, we exposed DMS-114 cells and JEB01 keratinocytes, derived from a junctional epidermolysis bullosa patient with a homozygous nonsense mutation (p.R688X) in the *COL17A1* gene, to increasing concentrations of G418 without or with pretreatment with 30 μM AC1903, the maximum dose tolerated by DMS-114 cells. Exposure of DMS-114 cells to G418 alone resulted in a concentration-dependent increase in full-length p53, which decreased ~60% when cells were additionally treated with AC1903 (**Figure 2A**). This reduction in PTC readthrough was correlated with significantly decreased intracellular G418 levels in the presence of AC1903 (**Figure 2B**). Similarly, JEB01 keratinocytes produced full-length Collagen XVII in the presence of G418 alone, which declined ~30-60% in combination with AC1903 (**Figure 2C**). Again, reduction of PTC readthrough in JEB01 cells exposed to AC1903 was correlated with significantly declined intracellular G418 levels (**Figure 2D**).

**Figure 2.**
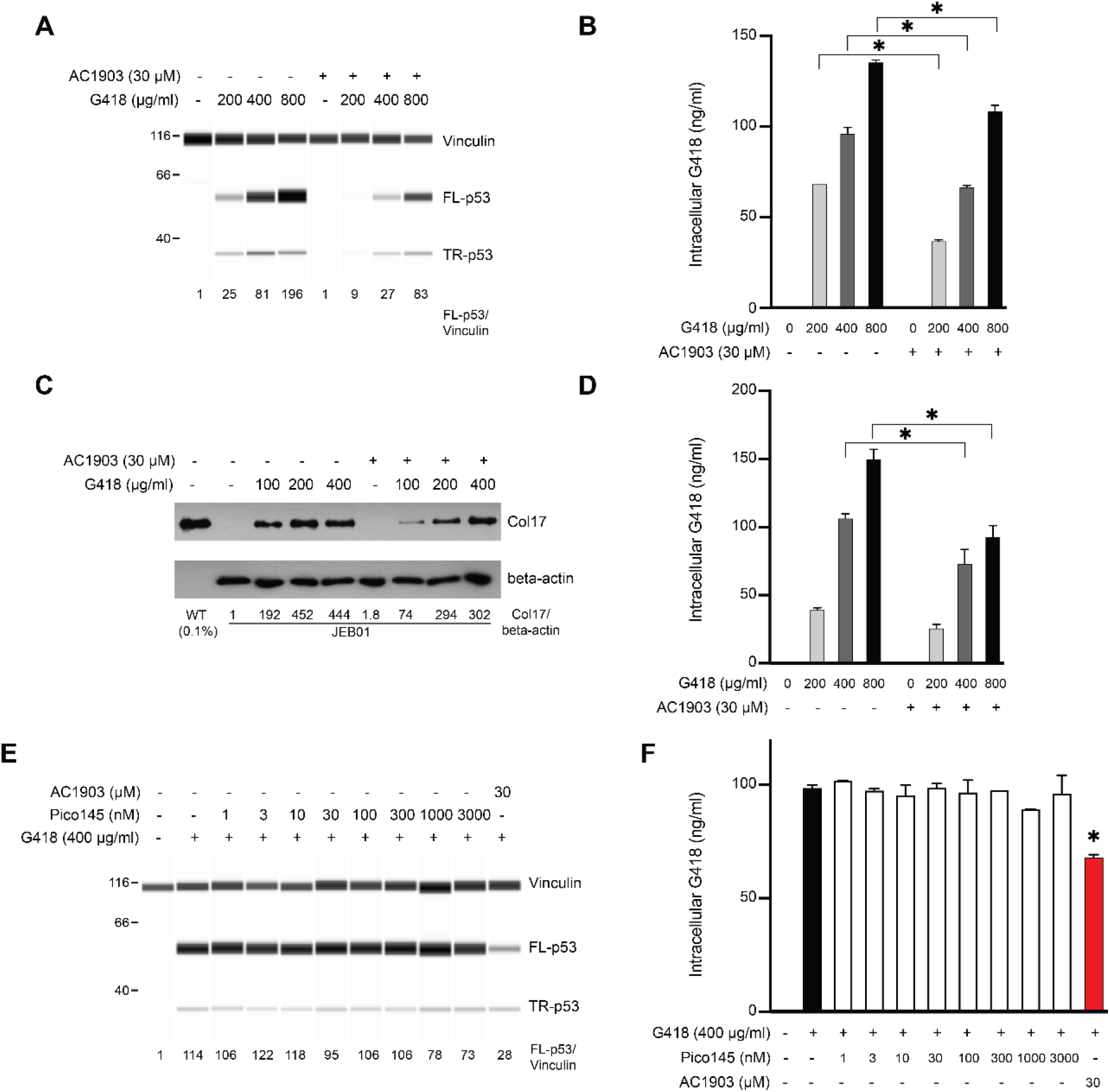
AC1903 suppresses G418 trafficking and PTC readthrough. A,C) DMS-114 (A) or JEB01 (C) cells were either left untreated or pretreated with 30 μM AC1903 for 3 h followed by exposure to indicated concentrations of G418 for another 3 h. At 24 h cell lysates were prepared and p53 (A) or collagen XVII (C) levels were measured. B,D) Total intracellular levels of G418 was measured in duplicates in the same lysates as in (A) and (C). E,F) DMS-114 cells were preincubated with 30 μM AC1903 or indicated concentrations of Pico145 for 3 h followed by exposure to 400 μg/ml G418 for another 3 h. At 24 h cell lysates were prepared and p53 (E) or intracellular G418 (F) levels were measured. In panels (A, C, E) protein levels were measured using automated capillary electrophoresis western analysis. For DMS-114 and JEB01 cell lysate analysis vinculin and beta-actin were used as loading controls, respectively. * indicates statistically significant difference between samples (P < 0.05).

These data suggest that AC1903 suppresses cellular uptake of G418 and G418-induced PTC readthrough in different cell lines, and the findings are consistent with a possible role of TRPC5 in cellular regulation of G418 uptake.

### G418 uptake and G418-induced PTC readthrough are not modulated by the TRPC1/4/5 inhibitor Pico145

In an attempt to validate the role of TRPC5 channels in the regulation of intracellular G418 levels by AC1903, we exposed DMS-114 cells to the xanthine derivative Pico145 in the presence of 400 μg/ml G418. Pico145 (also named HC-608) is a well-characterised, highly selective chemical probe of TRPC5, TRPC4 and heteromeric TRPC1/4/5 channels with picomolar to nanomolar potency (37, 43–45) that has been used successfully to inhibit endogenous TRPC1/4/5 channels in cells (43, 46–49), tissues (47, 50, 51) and animals (44, 51, 52). Unlike AC1903, Pico145 (up to 3 μM) did not affect either PTC readthrough or intracellular G418 levels (**Figure 2E–F**). These findings suggest that TRPC1/4/5 inhibition is not sufficient to mimic AC1903 as a suppressor of G418-induced PTC readthrough in DMS-114 cells.

### AC1903 is a non-selective TRP channel inhibitor

Based on the findings described above, we wondered whether AC1903 might have limited selectivity for TRPC5 channels and whether its effect on intracellular G418 levels and PTC readthrough might be the consequence of inhibition of other or multiple targets. All well-characterised TRPC channel inhibitors reported to date modulate multiple TRP(C) channels (32–34). Indeed, screening data deposited in PubChem by the John Hopkins Ion Channel Center suggest that AC1903 can also inhibit TRPC4 channels (IC_50_ 0.6 μM; PubChem Identifiers https://pubchem.ncbi.nlm.nih.gov/bioassay/434942#sid=865888AID434942-SID865888). Because limited data for the selectivity of AC1903 are available in the public domain, we decided to profile AC1903 against a selection of calcium-permeable cation channels by intracellular calcium ([Ca^2+^]_i_) recordings using well-characterised cell lines and ion channel activators (43, 53, 54), and the Ca^2+^ indicator Fura-2. In these assays, cells (HEK 293, HEK T-REx or CHO) over-expressing the relevant ion channel protein and loaded with Fura-2 AM (a cell-permeable precursor of Fura-2) were pre-incubated with various concentrations of AC1903 or vehicle (DMSO) for 30 minutes. Then, changes in ratiometric fluorescence (from background) were recorded upon addition of the respective ion channel activator.

We first established that our sample of AC1903 was analytically pure (**Figures S1–S4**) and that AC1903 does not interfere with the optical properties of Fura-2 (**Figures S5**). Then, we confirmed that AC1903 inhibits TRPC5:C5 channels activated by 30 nM EA (IC_50_ 18 μM) (**Figure 3A–B**) or 10 μM S1P, a physiological TRPC4/5 agonist (IC_50_ 3.5 μM) (**Figure S6A–B**). To test the effect of AC1903 on TRPC4 channels, we initially attempted to use carbachol to activate TRPC4:C4 channels, in line with patch-clamp experiments reported by Zhou et al. (41). However, application of carbachol also resulted in an increase in [Ca^2+^]_i_ in wild-type HEK 293 cells that lacked a response to EA, and AC1903 inhibited this carbachol-mediated calcium response (IC_50_ 20 μM) (**Figure S7**). Therefore, we used the selective TRPC1/4/5 activator EA and the physiological TRPC4/5 activator S1P instead. Consistent with data available on PubChem (see above), AC1903 inhibited TRPC4:C4 channels activated by 30 nM EA (IC_50_ 2.1 μM) (**Figure 3C–D**) or 10 μM S1P (IC_50_ 1.8 μM) (**Figure S6C–D**). Concentration-dependent inhibition of EA-activated TRPC4:C4 channels by AC1903 was subsequently confirmed by whole-cell patch-clamp recordings in HEK T-REx cells (**Figure S8**). Furthermore, AC1903 also inhibited EA-induced Ca^2+^ influx mediated by channels formed by concatemeric TRPC5–C1 (IC_50_ 4.7 μM) (**Figure S9A–B**) and TRPC4–C1 (IC_50_ 3.0 μM) (**Figure S9C–D**) (55–58), suggesting that AC1903 non-selectively inhibits TRPC1/4/5 channels. AC1903 also inhibited EA-mediated Ca^2+^ influx in A498 cells, which express endogenous TRPC4:C1 channels (IC_50_ 3.4 μM) (**Figure S9E–F**). AC1903 on its own did not affect viability of A498 cells (**Figure S10A**), but – consistent with inhibition of TRPC1:C4 channels (55) – AC1903 (at 100 μM) inhibited EA-mediated A498 cell death (**Figure S10B**).

**Figure 3.**
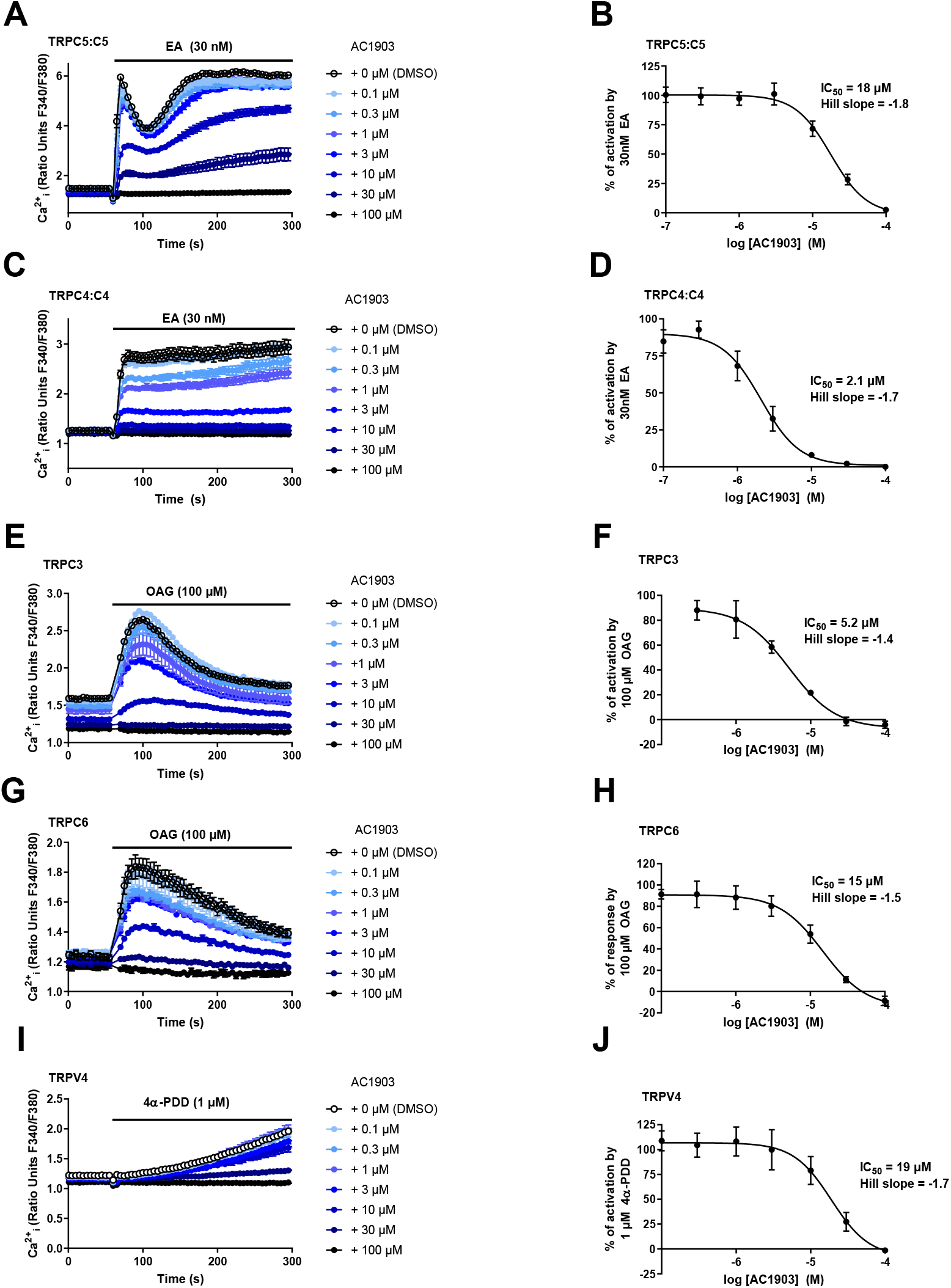
AC1903 inhibits multiple TRP channels. A,C) Representative [Ca^2+^]_i_ measurements from a single 96-well plate (N = 6) showing inhibition of EA-mediated [Ca^2+^]_i_ responses by 0.1-100 μM AC1903 in (Tet+) HEK T-REx cells expressing TRPC5 (A) or TRPC4 (C). B,D) Concentration-response data for experiments in (A) and (C), showing mean responses ± SEM (n/N = 3/18). Responses were calculated at 250-295 s compared to [Ca^2+^]_i_ at baseline (0-55 s). E,G) Representative [Ca^2+^]_i_ measurements from a single 96-well plate (N = 6) showing inhibition of OAG-mediated [Ca^2+^]_i_ responses by 0.1-100 μM AC1903 in (Tet+) HEK T-REx cells expressing TRPC3 (E) and WT HEK 293 cells transiently expressing TRPC6 (G). F, H) Concentration-response data for experiments in (E) and (G), showing mean responses ± SEM (n/N = 3/18). Responses were calculated at 90-110 s compared to [Ca^2+^]_i_ at baseline (0-55 s). (I) Representative [Ca^2+^]_i_ measurements from a single 96-well plate (N = 6) showing inhibition of the 4α-PDD-mediated [Ca^2+^]_i_ response by 0.1-100 μM AC1903 in CHO cells stably expressing TRPV4. J) Concentration-response data for experiments in (I), showing mean responses ± SEM (n/N = 3/18). Responses were calculated at 290-300 s compared to [Ca^2+^]_i_ baseline (0-55 s).

Further profiling studies showed that AC1903 can also inhibit OAG-activated channels formed by TRPC3 (IC_50_ 5.2 μM) (**Figure 3E–F**) and TRPC6 (IC_50_ 15 μM) (**Figure 3G–H**), as well as 4α-PDD-activated TRPV4 channels (IC_50_ 19 μM) (**Figure 3I–J**). In contrast, concentration of up to 30 μM of AC1903 did not inhibit capsaicin-activated TRPV1 or Yoda1-activated Piezo1 channels, but small effects were seen at 100 μM AC1903 (**Figure S11**). The combined data (summarised in **Table 1**) suggest that AC1903 can inhibit a range of TRP channels with similar (low micromolar) potencies, but that TRPV1 is relatively resistant.

**Table 1:** Summary of selectivity profiling of AC1903.

| **Table 1: Summary of selectivity profiling of AC1903** |  |  |  |
| --- | --- | --- | --- |
| **Ion channel (cell type)** | **Activator (concentration)** | **IC<sub>50</sub> (comment)** | **Data figures** |
| TRPC5:C5 (HEK T-REx) | EA (30 nM) | 18 μM | Figure 3A,B |
| TRPC5:C5 (HEK T-REx) | S1P (10 μM) | 3.5 μM | Figure S6A,B |
| TRPC5–C1 (HEK T-REx) | EA (30 nM) | 4.7 μM | Figure S9A,B |
| TRPC4:C4 (HEK T-REx) | EA (30 nM) | 2.1 μM<br>(consistent with patch-clamp data) | Figure 3C,D<br>Figure S8 |
| TRPC4:C4 (HEK T-REx) | S1P (10 μM) | 1.8 μM | Figure S6C,D |
| TRPC4–C1 (HEK T-REx) | EA (30 nM) | 3.0 μM | Figure S9C,D |
| TRPC1:C4 (A498) | EA (100 nM) | 3.4 μM<br>(inhibition of EA toxicity) | Figure S9E,F<br>Figure S10 |
| TRPC3:C3 (HEK T-REx) | OAG (100 μM) | 5.2 μM | Figure 3E,F |
| TRPC6:C6 (HEK 293) | OAG (100 μM) | 15 μM | Figure 3G,H |
| TRPV1:V1 (HEK 293) | capsaicin (500 nM) | >50 μM | Figure S11A,B |
| TRPV4:V4 (CHO) | 4α-PDD | 19 μM | Figure 3I,J |
| Piezo1 (HEK T-REx) | Yoda1 (3 μM) | >100 μM | Figure S11C,D |
| WT (HEK 293) | carbachol (100 μM) | 20 μM | Figure S7 |

## Discussion

In this study, we identified a suppressor of cellular uptake of the aminoglycoside G418 and G418-induced PTC readthrough. We initially observed a remarkable difference in G418-induced readthrough of the *TP53* gene between DMS-114 and TC-71 cells, despite both cell lines having the same nonsense mutation in *TP53*. We also found slightly lower uptake of G418 in TC-71 cells compared to DMS-114 cells. Because we identified a missense mutation in *TRPC5* in the TC-71 cell line, and because non-selective cation channels (including TRP channels) have been implicated in cellular uptake of aminoglycosides, we investigated the effect of the 2-aminobenzimidazole derivative AC1903, recently reported as a selective TRPC5 inhibitor. Indeed, AC1903 (10-30 μM) inhibited both G418 uptake and subsequent PTC readthrough in the DMS-114 cell line, as well as in JEB01 keratinocytes derived from a junctional epidermolysis bullosa patient. However, this effect could not be reproduced with the highly selective, nanomolar TRPC1/4/5 inhibitor Pico145 (using concentrations of up to 3 μM). These experiments suggested that TRPC1/4/5 inhibition is not sufficient to suppress G418 import, and that AC1903 may have previously unrecognised targets. Indeed, a PubChem search suggested that AC1903 can also inhibit TRPC4 channels (PubChem Identifiers https://pubchem.ncbi.nlm.nih.gov/bioassay/434942#sid=865888AID434942-SID865888). This prompted us to profile AC1903 against a range of non-selective cation channels.

Our experiments confirmed that AC1903 can inhibit calcium entry mediated by TRPC5:C5 channels (activated with EA or S1P), as well as EA-activated concatemeric TRPC5–C1 channels, which have functional and pharmacological properties similar to endogenous TRPC1:C5 channels. However, in contrast to the claim that AC1903 is a selective TRPC5 inhibitor with respect to TRPC4 and TRPC6 channels (41, 42), our data suggest that AC1903 can also inhibit calcium entry mediated by TRPC4:C4, TRPC4–C1, TRPC3:C3 and TRPC6:C6 channels, with IC_50_ values similar to or lower than those for TRPC5 channels. Moreover, AC1903 inhibited TRPV4 channels, but had limited effects on TRPV1 channels or Piezo1 channels. Discrepancies between some of our results and earlier reports on AC1903 could be related to differences between experimental protocols. For example, we measured the slow accumulation of [Ca^2+^]_i_ whereas Zhou et al. performed rapid measurements of currents, which include significant monovalent cation fluxes not detected in [Ca^2+^]_i_ recordings. In addition, Zhou et al. measured *acute* inhibition of conductivity of TRPC4, TRPC5 and TRPC6 channels by AC1903. Channels were activated with riluzole (TRPC5), carbachol (TRPC4) and OAG (TRPC6), respectively, and effects of AC1903 on activated channels were measured for ~ 20 s before AC1903 was washed out again. In contrast, we pre-incubated cells expressing the relevant TRPC channels with AC1903 for 30 min prior to addition of the relevant activator (EA or S1P for TRPC4:C4 and TRPC5:C5; EA for TRPC4–C1 and TRPC5–C1; OAG for TRPC3:C3 and TRPC6:C6; 4α-PDD for TRPV4:V4; Yoda1 for Piezo1). To address these differences, we performed whole-cell patch-clamp recordings, which showed acute, concentration-dependent inhibition of EA-activated TRPC4:C4 channels by AC1903. It is possible that AC1903-mediated inhibition of TRPC6 channels is slower than inhibition of TRPC4 and TRPC5 channels, which may explain that this effect was not be detected during the short application protocol used by Zhou et al. Effects of AC1903 on different TRPC channels may also be channel state- or voltage-dependent.

High-quality chemical probes are powerful tools in target validation studies, and can serve as useful starting points for drug development, especially when combined with genetic approaches. In our recent review, we highlighted the use of small-molecule modulators of TRPC1/4/5 channels to complement genetic approaches in dissecting the different roles of specific TRPC1/4/5 channels across species, tissues, and pathologies (34). However, even high-quality chemical probes are likely to have off-targets, and potency and selectivity of a chemical probe may be dependent on factors such as cellular context, mode of application, and treatment time. As part of the development of AC1903, the compound was screened against a standard kinase panel, in which no off-target effects were found (41). However, our data suggest that TRPC3, TRPC4, TRPC6 and TRPV4 are important potential off-targets of AC1903. In addition, the inhibition by AC1903 of the carbachol-mediated calcium response in wild-type HEK 293 cells (which lacked response to EA) suggests that AC1903 targets additional cellular calcium handling mechanisms. Therefore, data from experiments in which AC1903 is used as a TRPC5 inhibitor need to be interpreted with these results in mind, and additional chemical and genetic approaches may be required to dissect the roles of specific channels.

Because selective inhibition of TRPC1/4/5 channels with Pico145 could not reproduce the effects of AC1903 in our experiments, it is possible that AC1903 suppresses PTC readthrough by inhibition of multiple TRP channels involved in G418 uptake. This hypothesis is consistent with our pharmacological profiling of AC1903 and with previous reports suggesting that multiple non-selective cation channels (e.g., TRPV1 and TRPV4) can mediate import of aminoglycosides (24, 25, 28). The potential for direct permeation of small organic cations through the pore of non-selective cation channels (e.g., P2X (59–62), TRPV1 (25, 28, 63–66), TRPV2 (67–69), TRPV4 (24, 28) and TRPA1 (70)) is a topic of multiple investigations (71). In addition, a recent cryo-EM structure of a TRPV2 channel showed that doxorubicin (an organic cation at physiological pH) can enter the TRPV2 pore in the presence of the TRPV2 activator 2-APB (72). Determination of specific TRP channels that may be involved in the suppression of G418 uptake by AC1903 is challenging. TRP proteins (especially those of the TRPC sub-family) can form various homo- and heterotetrameric channels with distinct biophysical properties, localisation, pharmacology and function, and genetic perturbation of TRP proteins can cause alterations of channel stoichiometries as well as compensatory upregulation of other (TRP) ion channels. In addition, it remains possible that AC1903 suppresses G418-induced PTC readthrough via other mechanisms. Therefore, further work is required to determine its mode-of-action.

The potential implication of the R175H mutation near the putative zinc binding site of TRPC5 – which also incorporates cysteines shown to undergo S-glutathionylation and S-palmitoylation – in suppression of G418-induced PTC readthrough is currently not understood. In addition, it is difficult to explain why a small difference in intracellular G418 concentrations between DMS-114 and TC-71 cells would result in such profound differences in formation of PTC readthrough product, suggesting that additional cellular mechanisms may be involved. Further work is required to determine if, and how, the R175H mutation affects TRPC5 channel localisation and function. This mutation of *TRPC5* is hemizygous in the TC-71 cell line, further increasing the number of potential homo- and heteromeric channels that could incorporate TRPC5.

In conclusion, our work reveals AC1903 as a suppressor of G418 uptake and G418-induced PTC readthrough, and suggests that further studies are required to understand the roles of various TRP channels in aminoglycoside uptake in different cell types. Such understanding may guide studies of PTC readthrough in cells and animal models to determine the therapeutic potential of aminoglycosides in the treatment of rare genetic diseases.

## Materials and Methods

### Human Cells for PTC readthrough

DMS-114 and TC-71 cell lines with homozygous nonsense mutation in the *TP53* gene (NM 000546.5:c.637C>T; NP 000537.3:p.R213X) were purchased from the American Type Culture Collection and the German Collection of Microorganisms and Cell Cultures, respectively. DMS-114 and TC-71 cells were cultured in RPMI-1640 medium (Sigma-Aldrich, ON, Canada) supplemented with 10% (vol/vol) FBS and 1% antibiotic-antimycotic at 37 °C and 5% (vol/vol) CO_2_. Immortalised JEB01 keratinocytes were derived from a junctional epidermolysis bullosa (JEB) patient with a homozygous nonsense mutation in the *COL17A1* gene (NM_000494.4:c.2062C > T;NP_000485.3:p.R688X) (73). JEB01 keratinocytes were cultured in defined keratinocyte serum-free medium (K-SFM) supplemented with defined K-SFM growth supplement (Gibco/Thermo Fisher Scientific, MD, USA) and 1% antibiotic– antimycotic (Gibco/Thermo Fisher Scientific, MD, USA) at 37 °C and 5% (vol/vol) CO_2_.

### Cell Lines for TRP Channel Assays and Viability Studies

HEK T-REx™ cells expressing tetracycline-inducible TRPC4, TRPC5, TRPC5–C1,TRPC4– C1 and hPiezo1 have been described previously (53, 55, 57, 74). Cloning of TRPC3 has been described previously.(75) To create cells stably expressing tetracycline-inducible TRPC3, HEK T-REx cells were transfected with pcDNA4/TO/hTRPC3 using jetPRIME ® transfection reagent (VWR, Lutterworth, UK) according to manufacturer’s instructions. After 48 h cells were put under antibiotic selection using 400 μg/ml zeocin and 10 μg/ml Blasticidin S (Invivogen, San Diego, California, US)). Medium changes were carried out every 2 to 3 days to remove dead cells. Inducible HEK T-REx cells were cultured in Dulbecco’s modified Eagle medium GlutaMAX (Thermo Fisher Scientific, Paisley, UK) containing 10% fetal bovine serum (FBS; Merck, Gillingham, UK), penicillin-streptomycin (100 units ml^−1^/ 100 μg ml^−1^), with the addition of blasticidin (10 μg ml^1^; Invivogen) and zeocin (Invivogen) at either 400 μg ml^−1^ (TRPC cell lines) or 200 μg ml^−1^ (hPiezo1) to maintain the stable incorporation of the tetracycline repressor and the channel of interest, respectively. Expression of tetracycline-inducible proteins was induced by the addition of tetracycline (TRPC cell lines: 1 μg ml^−1^, hPiezo1: 100 ng ml^−1^; 24 hr) to cell culture medium. Experiments with TRPC6 and TRPV1 were carried out in WT HEK 293 cells transiently transfected with hTRPC6 (54) or rat TRPV1 (plasmid donated by Prof. Nikita Gamper, University of Leeds). WT HEK 293 cells were maintained in DMEM containing 10% FBS and penicillin-streptomycin (100 units ml^−1^ / 100 μg ml^−1^). Cells were transfected with hTRPC6 (cloned into pcDNA3) or rat TRPV1 using jetPRIME ® transfection reagent (VWR, Lutterworth, UK). Cells were assayed 48 hours after transfection. CHO cells stably expressing TRPV4 (43) were cultured in Ham’s Nutrient Mixture F12 (Thermo Fisher Scientific), supplemented with 10% FBS, penicillin-streptomycin (100 units ml^−^/ 100 μg ml^−1^), and 1 mg ml^−1^ G418 (Invivogen) to maintain expression of TRPV4. A498 cells were cultured in Minimum Essential Medium (MEM) with Earle’s salts (Thermo Fisher Scientific) supplemented with 10 % fetal bovine serum and penicillin-streptomycin (100 units ml^−1^/ 100 μg ml^−1^).

### Automated Capillary Electrophoresis Western Analysis

The western analysis assays for p53 and Vinculin detection were performed as previously described (18). Briefly, mixtures of 1 mg ml^−1^cell lysates and the fluorescent master mix were heated at 95 °C for 5 min. The samples and all other reagents were dispensed into the microplates and capillary electrophoresis western analysis was carried out with the ProteinSimple WES instrument and analysed using the inbuilt Compass software (ProteinSimple, CA, USA). DO-1 mouse anti-p53 antibody (1:400, Santa Cruz sc-126, TX, USA) and mouse anti-vinculin antibody (1:600, R&D Systems MAB6896, MN, USA) were used.

### SDS-PAGE and immunoblotting

Human JEB01 keratinocytes were seeded into 6-well tissue culture dishes and exposed to indicated compounds. After 24 h, cells were lysed and 15 μg total protein from each lysate was separated on a 6% polyacrylamide gel, electrotransferred onto a nitrocellulose membrane, blocked, incubated with rabbit anti-Collagen XVII antibody (1:1,000, Abcam ab184996, ON, Canada) overnight at 4 °C, washed and incubated with HRP-conjugated goat anti-rabbit secondary antibody and developed using enhanced chemiluminescence substrate (Millipore, ON, Canada). Membrane was stripped using 0.1 N NaOH and reprobed with rabbit anti-beta actin antibody (1:10,000, Novus Biologicals, CO, USA) and detected as above.

### Measurement of intracellular G418

Intracellular G418 was measured in cell lysates prepared for western analysis using an indirect competitive gentamicin ELISA kit (Creative Diagnostics, DEIA047, NY, USA), according to the manufacturer’s protocol, as previously described (76). Standard curves were generated using G418 diluted in lysis buffer.

### Intracellular Ca^2+^ measurements

[Ca^2+^]_i_ recordings were carried out using the ratiometric Ca^2+^ dye Fura-2. 24 hours prior to experiments, cells were plated either onto black, clear bottom poly-D-lysine coated 96 well plates (HEK cells) or clear 96 well plates (A498 cells, CHO cells) at 50,000 cells per well (HEK cells) or 20,000 cells per well (A498 cells, CHO cells). For tetracycline-inducible HEK T-REx cells, protein expression was induced with tetracycline (TRPC channels: 1 μg ml^−1^; hPiezo1: 100 ng ml^−1^) at this point. To load cells with the Fura-2 dye, media was removed and cells were incubated with standard bath solution (SBS) containing 2 μM Fura-2 acetoxymethyl ester (Fura-2 AM; Thermo Fisher Scientific, Paisley, UK) and 0.01% pluronic acid for 1 h at 37 °C. SBS contained (in mM): NaCl 130, KCl 5, glucose 8, HEPES 10, MgCl_2_ 1.2 and CaCl_2_ 1.5. After this incubation, cells were washed twice with fresh SBS. SBS was then changed to recording buffer, which consisted of SBS with 0.01% pluronic acid and the relevant concentration of AC1903, diluted in DMSO (final DMSO concentration: 0.01%). Control wells contained SBS with 0.01% pluronic acid and 0.01% DMSO. Cells were then incubated for 30 min prior to the experiment. For experiments with CHO cells, probenecid at 2.5 mM was included throughout the experiment to prevent extrusion of Fura-2 from the cells. Measurements were carried out using a FlexStation (Molecular Devices, San Jose, CA), using excitation wavelengths of 340 and 380 nm, at an emission wavelength of 510 nm. [Ca^2+^]_i_ recordings were performed at room temperature at 5 s intervals for 300 s. Compounds were added from a compound plate at 2x the final concentration after recording for 60 s.

### Whole-cell patch-clamp recordings

For electrophysiology experiments, cells were plated at a low density of 20-30% onto round coverslips (13 mm diameter or 5 × 5 mm), and TRPC4 expression in HEK T‐REx was induced with 1 μg·ml^−1^ tetracycline 24 h before experimentation. Experiments were carried out at room temperature. Patch‐clamp recordings were performed in whole‐cell mode under voltage clamp at room temperature using 3-5 MΩ patch pipettes fabricated from borosilicate glass capillaries with an outside diameter of 1 mm and an inside diameter of 0.58 mm (Harvard Apparatus). The voltage protocol comprised voltage ramps applied from −100 to +100 mV every 10 s from a holding potential of 0 mV. The patch‐clamp currents were recorded using a Axopatch 200B amplifier, digitised by a Digidata 1440 and recorded to a computer using pCLAMP10 (Molecular Devices). The data were filtered at 1 kHz and analysed offline using Clampfit 10.7 software and Origin 2019b software (OriginLab, Northampton, MA). The bath solution consisted of SBS and the pipette solution (intracellular solution) contained (in mM) CsCl 145, MgCl_2_ 2, HEPES 10, EGTA (free acid) 1, ATP (sodium salt) 5, NaGTP 0.1, titrated to pH 7.2 with CsOH. All solutions were filtered using a 0.2 μm filter (Nalgene Rapid Flow, Thermo Scientific). Upon whole cell configuration, cells were continually perfused at a rate of 2 ml·ml^−1^ with SBS containing 0.01% pluronic acid, followed by SBS + 0.01% pluronic acid + EA (30 nM) (**Figure S8**).

### Cell Viability Assay

Cell viability assays were carried out using WST-1 Cell Proliferation Reagent (Merck). A498 cells were plated onto clear 96-well plates at 4,000 cells per well and left to adhere for 24 hours. Media was then removed, and replaced with fresh media containing 100 nM EA, in presence of either DMSO or 0.1 – 100 μM AC1903. Control wells were treated with vehicle (DMSO) only. To exclude any potential cytotoxic effects of AC1903 itself, cells were incubated with 0.1 – 100 μM AC1903 in absence of EA. Cells were treated in triplicate. Cells were incubated with compounds for 8 hours. After 8 hours, treatment media was removed and media containing WST-1 reagent (10 μL reagent + 100 μL media per well) was added to each well and incubated for 1.5 hours at 37 °C. Blank wells contained media and WST-1 reagent (no cells). After incubation, the plate was shaken for 1 minute. Absorbance was then read at 440 nm and 650 nm (reference wavelength). To analyse data, the value of the 650 nm was subtracted from that of the 440 nm reading. Values across triplicate wells were averaged, and the value from the blank wells subtracted to account for background absorbance. Values from compound -treated cells were then compared to control cells (DMSO-treated).

### Data Analysis

For G418 uptake experiments, statistical analysis was performed using GraphPad Prism 8.0. Two-way Analysis of Variance (ANOVA) was used to analyse the difference between different treatments or samples and differences were considered significant at a P-value of < 0.05.

For Ca^2+^ recording and electrophysiology experiments, data were analysed using GraphPad Prism 9.0. Representative data are presented as raw data. Unless indicated otherwise, experiment were carried out at n = 3, where n = number of independent experiments. Each independent FlexStation experiment consisted of 6 technical replicates (i.e. 6 wells of a 96 well plate for each Flex Station assay), unless stated otherwise. Concentration-response curves were fitted in GraphPad Prism using a four parameter curve fit. Amplitudes of Ca^2+^ responses for different channels were measured at timepoints indicated in the corresponding figure legends.

### Absorbance and Fluorescent Properties of AC1903

AC1903 or vehicle (DMSO) was dissolved in SBS to the stated concentration and added to wells of a clear 96 well plate. Spectra were recorded using a FlexStation. Absorbance was recorded from 200-850 nm at 10 nm intervals. A fluorescence excitation spectrum was recorded from 250-800 nm at 10 nm intervals with emission fixed at 510 nm (the emission wavelength of Fura-2). Fluorescence emission spectra were recorded from 250-800 nm at 10 nm intervals, with excitation fixed at either 340 nm or 380 nm (excitation wavelengths for Fura-2).

### Chemicals

(-)-Englerin A (EA) was obtained from PhytoLab (Vestenbergsgreuth, Germany), AC1903 was obtained from Cayman Chemical (Ann Arbor, USA) and identity and purity were confirmed through analysis by ^1^H NMR, ^13^C NMR, high-resolution mass spectrometry and HPLC analysis (**Figures S8–S11**); data were consistent with previous reports (42). 1-oleoyl-2-acetyl-sn-glycerol (OAG) and 4α-phorbol 12,13-didecanoate (4α-PDD) were obtained from Merck (Gillingham, UK). Sphingosine-1-phosphate (S1P) and capsaicin were obtained from Bio-techne (Abingdon, UK). Carbachol was obtained from Alfa Aesar Chemicals (Heysham, UK). Yoda1 was obtained from Tocris Bioscience (Bristol, UK). Chemicals were made up as 10 mM (EA, 4α-PDD, Yoda-1) or 100 mM (AC1903, OAG) stocks in 100% DMSO, aliquots of which were stored at −20 °C (Yoda1, AC1903) or −80 °C (EA, 4α-PDD, OAG)). S1P was made up in methanol to 5 mM and stored as aliquots at −80 ⁰C. Further dilutions of EA and AC1903 were made in DMSO if required and these were dissolved 1:1000 in recording and compound buffer (SBS + 0.01% pluronic acid) before being added to cells. Fura-2-AM (Thermo Fisher Scientific) was dissolved at 1 mM in DMSO.

## Abbreviations

TRPC: Transient Receptor Potential Canonical
EA: (-)-englerin A
S1P: sphingosine-1-phosphate
OAG: 1-oleoyl-2-acetyl-sn-glycerol
HEK T-REx cells: human embryonic kidney 293 cells stably expressing the tetracycline repressor protein, allowing tetracycline-inducible recombinant protein over-expression
PTC: premature termination codon
2-APB: 2-aminoethoxydiphenylborane
4α-PDD: 4α-phorbol 12,13-didecanoate
TRPC1, TRPC3, TRPC4, TRPC5: TRPV1 and TRPV4 denote the different proteins or channels incorporating them
TRPC1/4/5 denotes channels composed of TRPC1: TRPC4 and/or TRPC5 (homo- or heteromeric; any ratio)
TRPC3:C3, TRPC4:C4, TRPC5:C5, TRPC6:C6: TRPV1:V1 and TRPV4:V4 denote specific homomeric channels
TRPC1:C4 and TRPC1:C5 denote heteromeric channels formed by TRPC1 and either TRPC4 or TRPC5 (any ratio): TRPC4–C1 and TRPC5–C1 denote (channels composed of) recombinant, concatemeric proteins (fusions of TRPC1 at the C-terminus of either TRPC4 or TRPC5 through a short linker)
[Ca^2+^]_i_: intracellular concentration of Ca^2+^

## Data availability

Data relating to this manuscript are available from the corresponding authors upon reasonable request.

## Acknowledgments

We thank Prof. Nikita Gamper for donation of a TRPV1 plasmid.

## Author contributions

ABH, CCB, IBP, SHF, ADB, KC and DML performed experiments. ABH, CCB, IBP, DML, SHF, ADB, DJB and RSB designed experiments and analysed data. ABH, CCB and IBP made figures. ABH and RSB conceived and led the project. MR, DJB and RSB generated research funds. ABH, CCB and RSB wrote the manuscript. All authors commented on the manuscript.

## Funding and additional information

This work was supported by an ERA-Net E-RARE grant funded by the Canadian Institute for Health Research (ERT-155725 to MR), the BBSRC (BB/P020208/1 to DJB and RSB), the BHF (PG/19/2/34084 to DJB and RSB), and a Wellcome Trust PhD studentship (102174/B/13/Z) to IBP.

---

**Figure S1.**
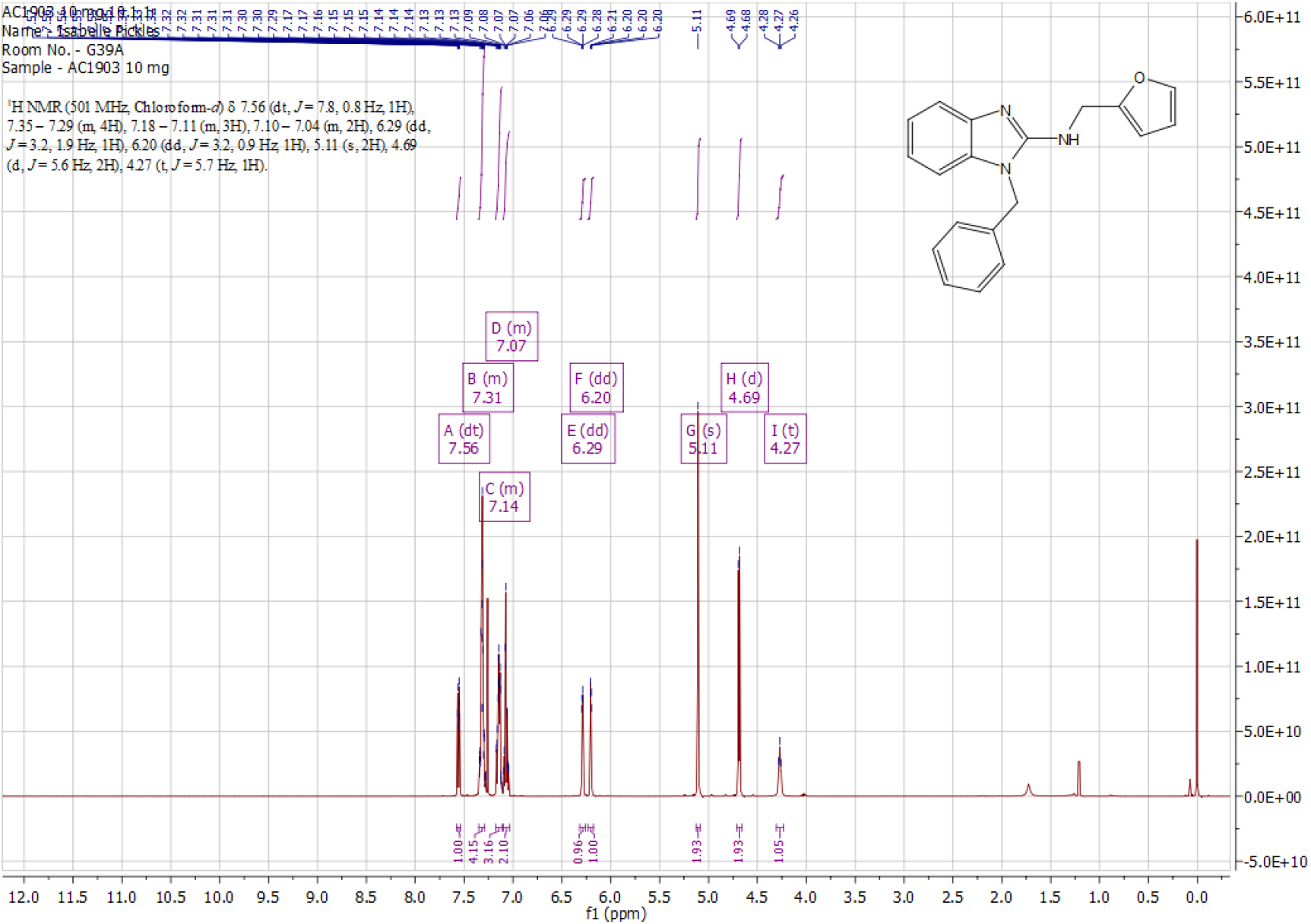
^1^H NMR spectrum of commercial AC1903 (Cayman Chemical, Ann Arbor, USA).

**Figure S2.**
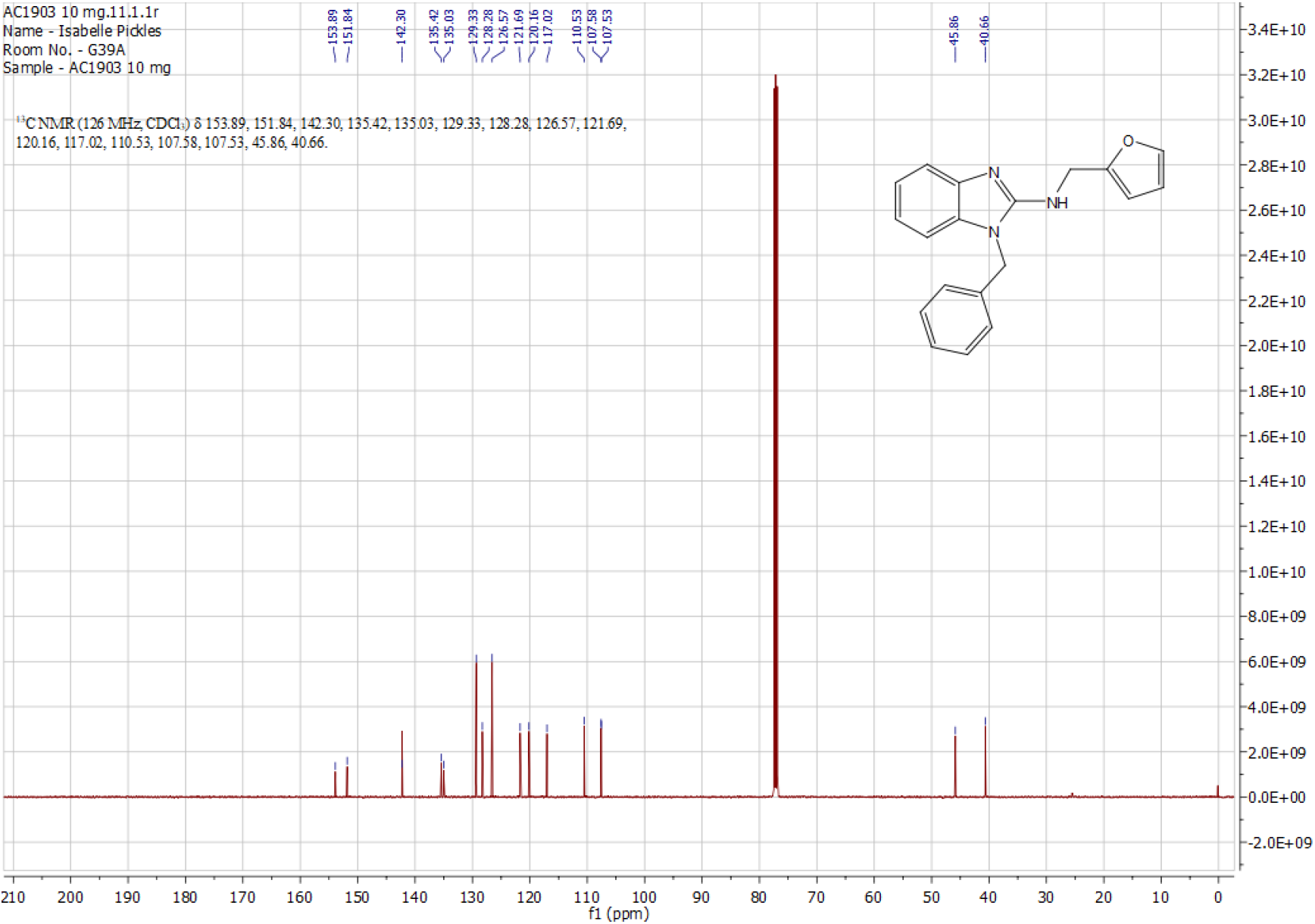
^13^C NMR spectrum of commercial AC1903 (Cayman Chemical, Ann Arbor, USA).

**Figure S3.**
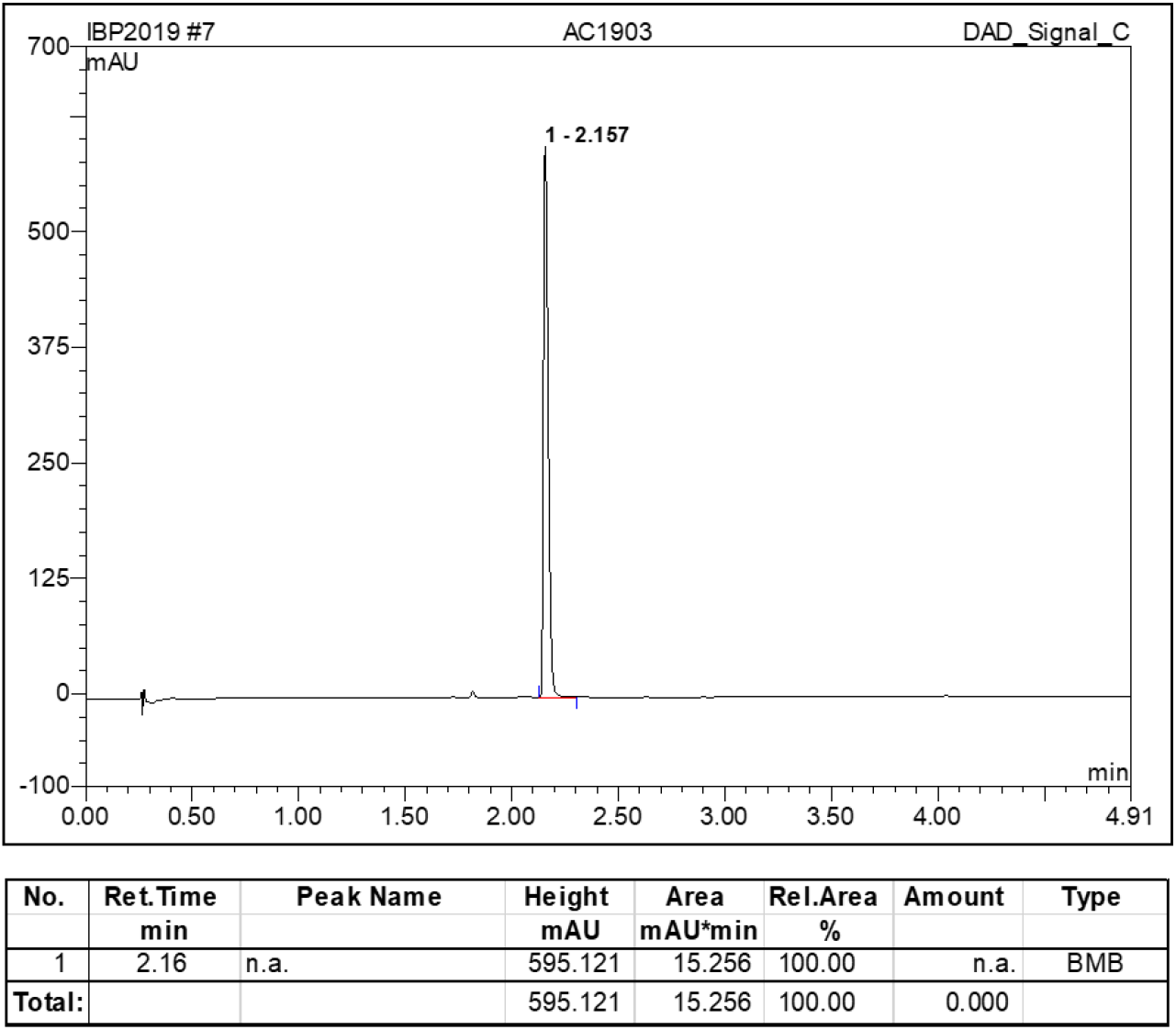
HPLC trace of commercial AC1903 (Cayman Chemical, Ann Arbor, USA). Analysis was performed on an Agilent 1290 Infinity Series equipped with a UV detector (set at 254 nm) and a Hyperclone C18 reverse phase column using MeCN/water (5→95% or 50→95%) containing 0.1% formic acid, at 0.5 mL min^−1^ over a period of five minutes.

**Figure S4.**
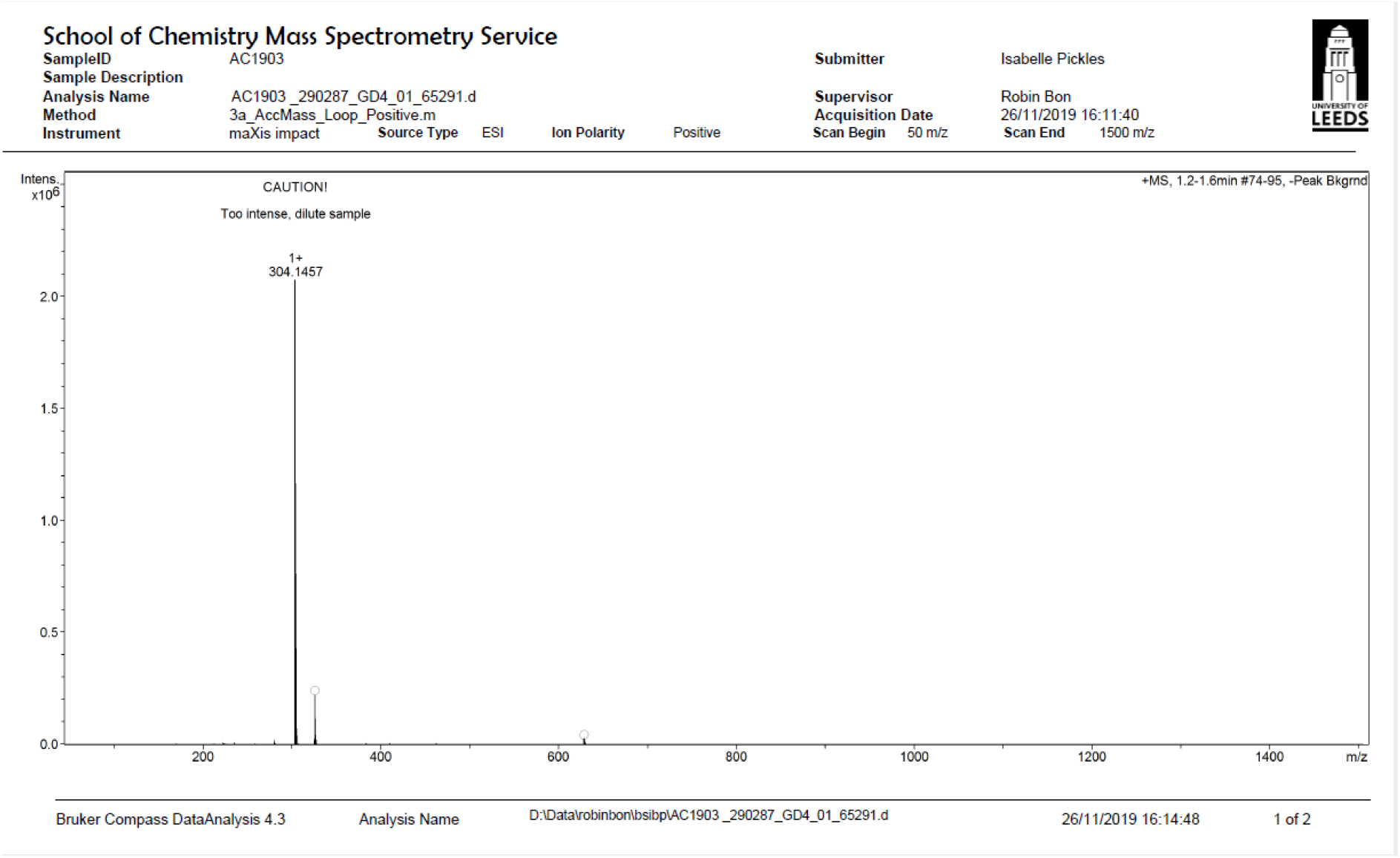
HRMS spectrum of commercial AC1903 (Cayman Chemical, Ann Arbor, USA). **1-Benzyl-N-(furan-2-ylmethyl)-1H-benzo[d]imidazol-2-amine, AC1903 (Cayman Chemical, Ann Arbor, USA)** ^1^H NMR (501 MHz, Chloroform-d) δ 7.56 (dt, J = 7.8, 0.8 Hz, 1H), 7.35 – 7.29 (m, 4H), 7.18 – 7.11 (m, 3H), 7.10 – 7.04 (m, 2H), 6.29 (dd, J = 3.2, 1.9 Hz, 1H), 6.20 (dd, J = 3.2, 0.9 Hz, 1H), 5.11 (s, 2H), 4.69 (d, J = 5.6 Hz, 2H), 4.27 (t, J = 5.7 Hz, 1H); ^13^C NMR (126 MHz, CDCl3) δ 153.9, 151.8, 142.3, 135.4, 135.0, 129.3, 128.3, 126.6, 121.7, 120.2, 117.0, 110.5, 107.6, 107.5, 45.9, 40.7; ESI-HRMS: calc. for C_19_H_17_N_3_NaO [M+Na]^+^ 326.1264, found 326.1263; ESI-LCMS: m/z 304.21 [M+H]+; HPLC (5-95 % MeCN in water) RT = 2.16 min.

**Figure S5.**
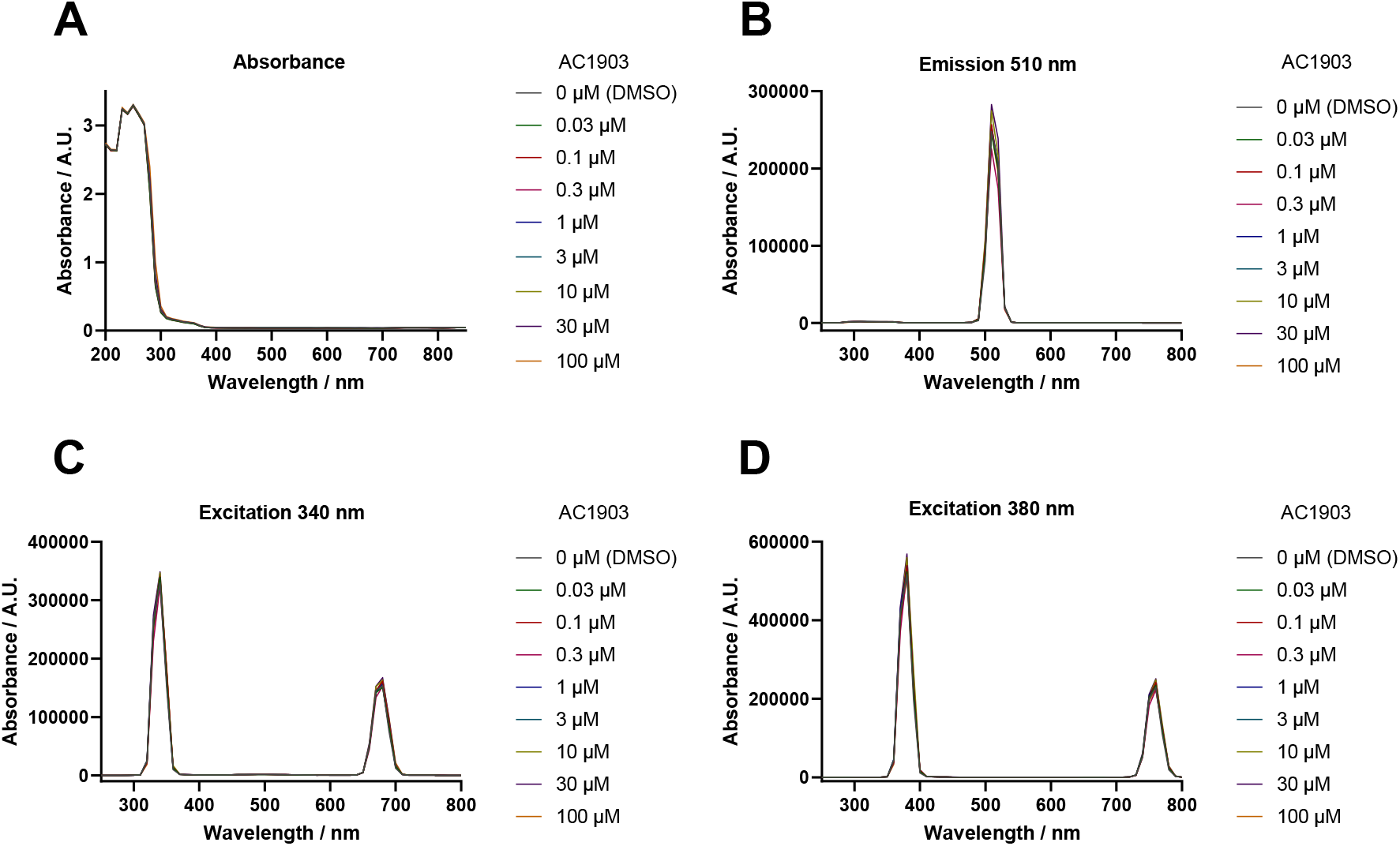
Absorbance and fluorescence spectra of solutions containing AC1903 (Cayman Chemical, Ann Arbor, USA). A) Absorbance spectra of SBS containing DMSO or 0.03-100 μM AC1903. Absorbance was measured at 200-850 nm at 10 nm intervals. B) Excitation spectra of SBS containing DMSO or 0.03-100 μM AC1903. Emission was fixed at 510 nm, with excitation of 250−800 nm at 10 nm intervals. C,D) Fluorescence emission spectra of SBS containing DMSO or 0.03-100 μM. Excitation wavelength was fixed at either 340 nm (C) or 380 nm (D), while emission was measured at 250−800 nm, at 10 nm intervals.

**Figure S6.**
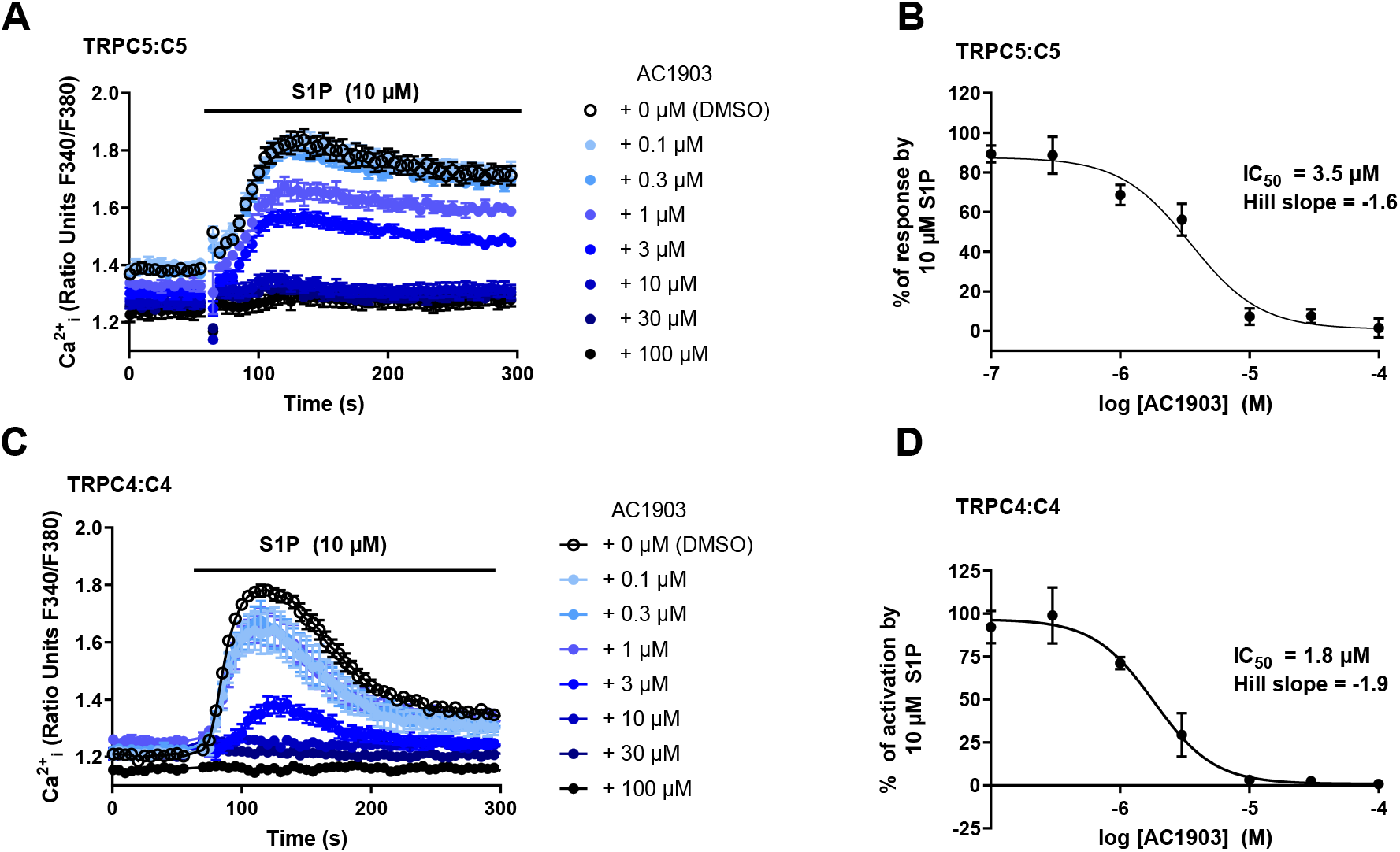
AC1903 inhibits S1P-mediated activation of TRPC5:C5 and TRPC4:C4 channels. A,C) Representative [Ca^2+^]_i_ measurements from a single 96-well plate (N = 6) showing inhibition of 10 μM S1P-mediated [Ca^2+^]_i_ responses by 0.1-100 μM AC1903 in (Tet+) HEK T-REx cells expressing TRPC5 (A) or TRPC4 (C). B,D) Concentration-response data for experiments in (A) and (C), respectively, showing mean responses ± SEM (n/N = 3/18). Responses were calculated at 100-150 s compared to [Ca^2+^]_i_ at baseline (0-55 s).

**Figure S7.**
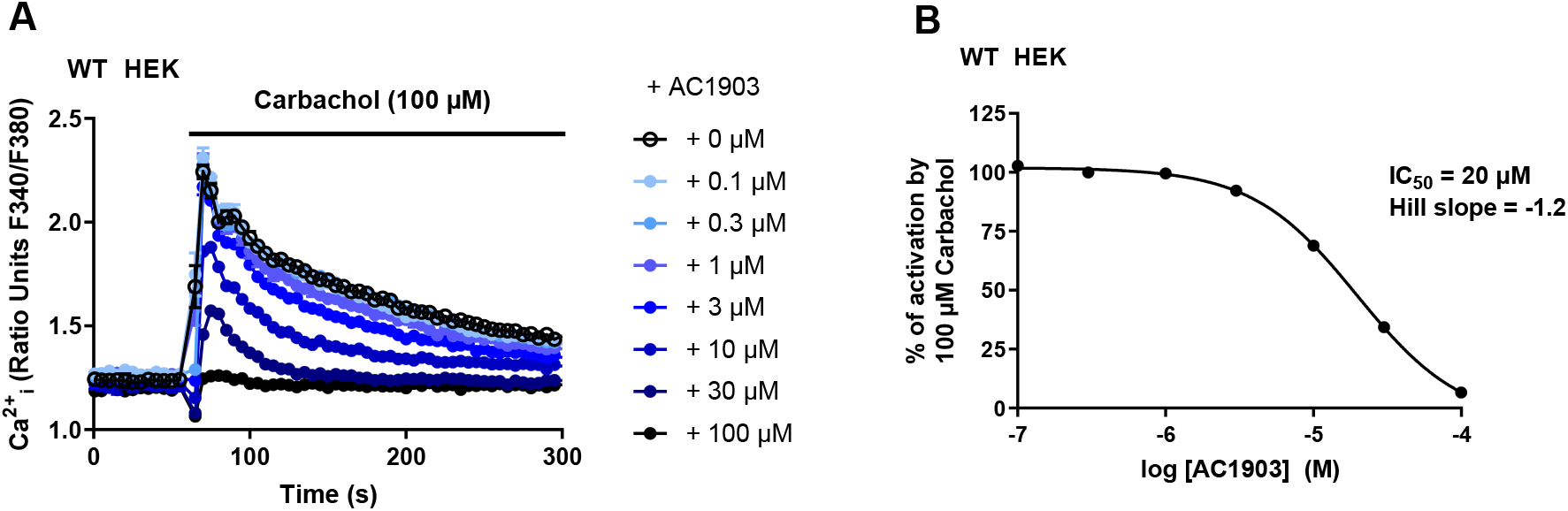
AC1903 inhibits carbachol-mediated [Ca^2+^]_i_ responses in WT HEK 293 cells. A) Representative [Ca^2+^]_i_ measurements from a single 96-well plate (N = 6) showing inhibition of carbachol-mediated [Ca^2+^]_i_ responses by 0.1-100 μM AC1903 in WT HEK 293 cells. B) Concentration-response data for experiments in (A), showing mean responses ± SEM (n/N = 1/6). Responses were calculated at 70−80 s compared to [Ca^2+^]_i_ at baseline (0-55 s).

**Figure S8.**
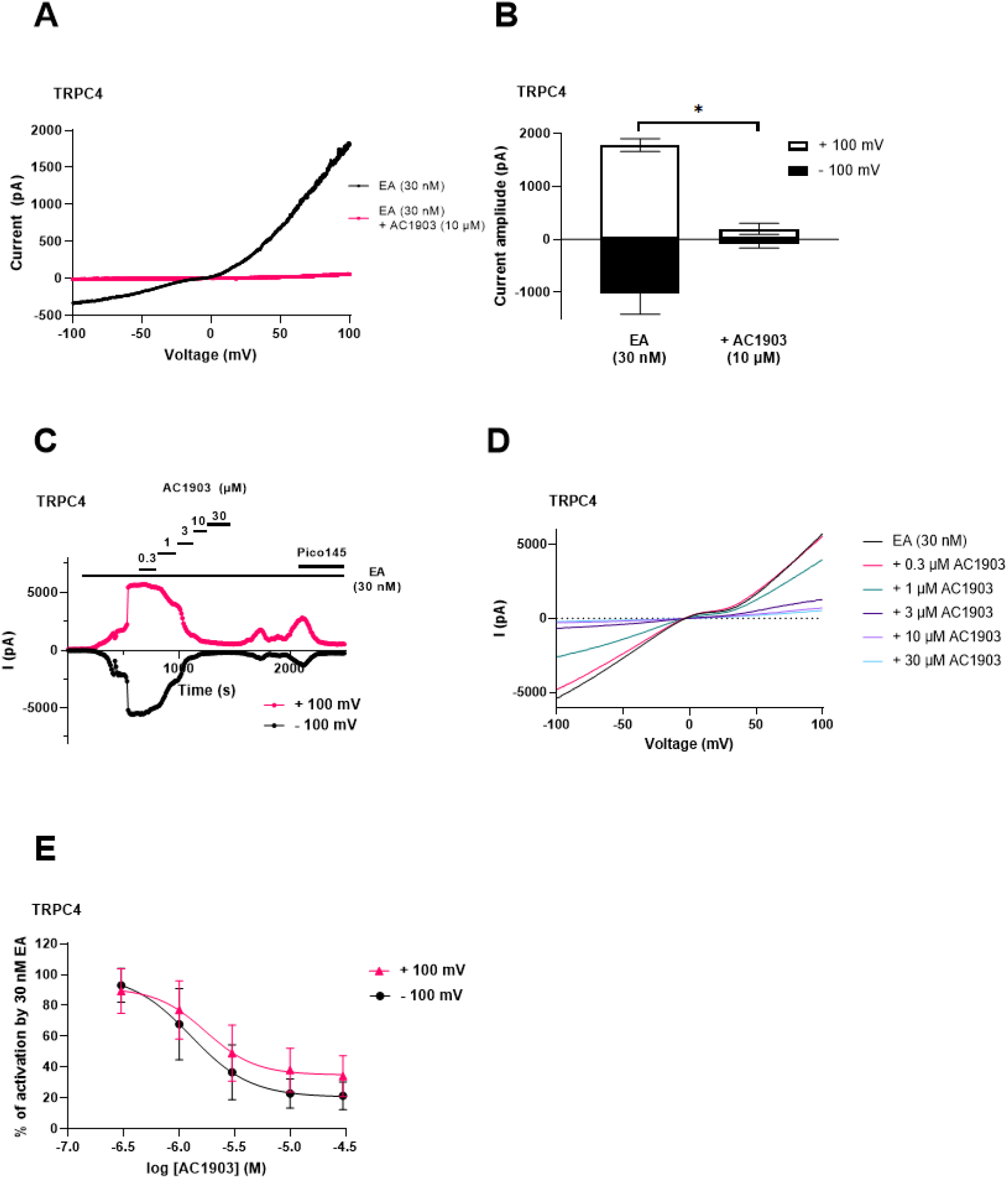
AC1903 inhibits TRPC4 activation in whole-cell patch-clamp recordings. A) Representative current-voltage relationship for one (Tet+) HEK T-Rex cell expressing TRPC4, showing current after activation by EA (30 nM) and subsequent inhibition by addition of AC1903 (10 μM). B) Quantified data for experiments in (A), showing mean ± SEM for current at +100 mV (white bars) or −100 mV (black bars) (n = 3). C) Representative trace from one (Tet+) HEK TREx cell expressing TRPC4, showing current at +100 mV (magenta) and −100 mV (black) after activation by EA (30 nM), followed by cumulative addition of AC1903 (0.3-30 μM). D) Current-voltage relationships for experiment in (C). E) Concentration-response data for experiments in (C), showing % of activation by 30 nM EA at +100 mV (magenta triangles) and −100 mV (black circles) (n = 3).

**Figure S9.**
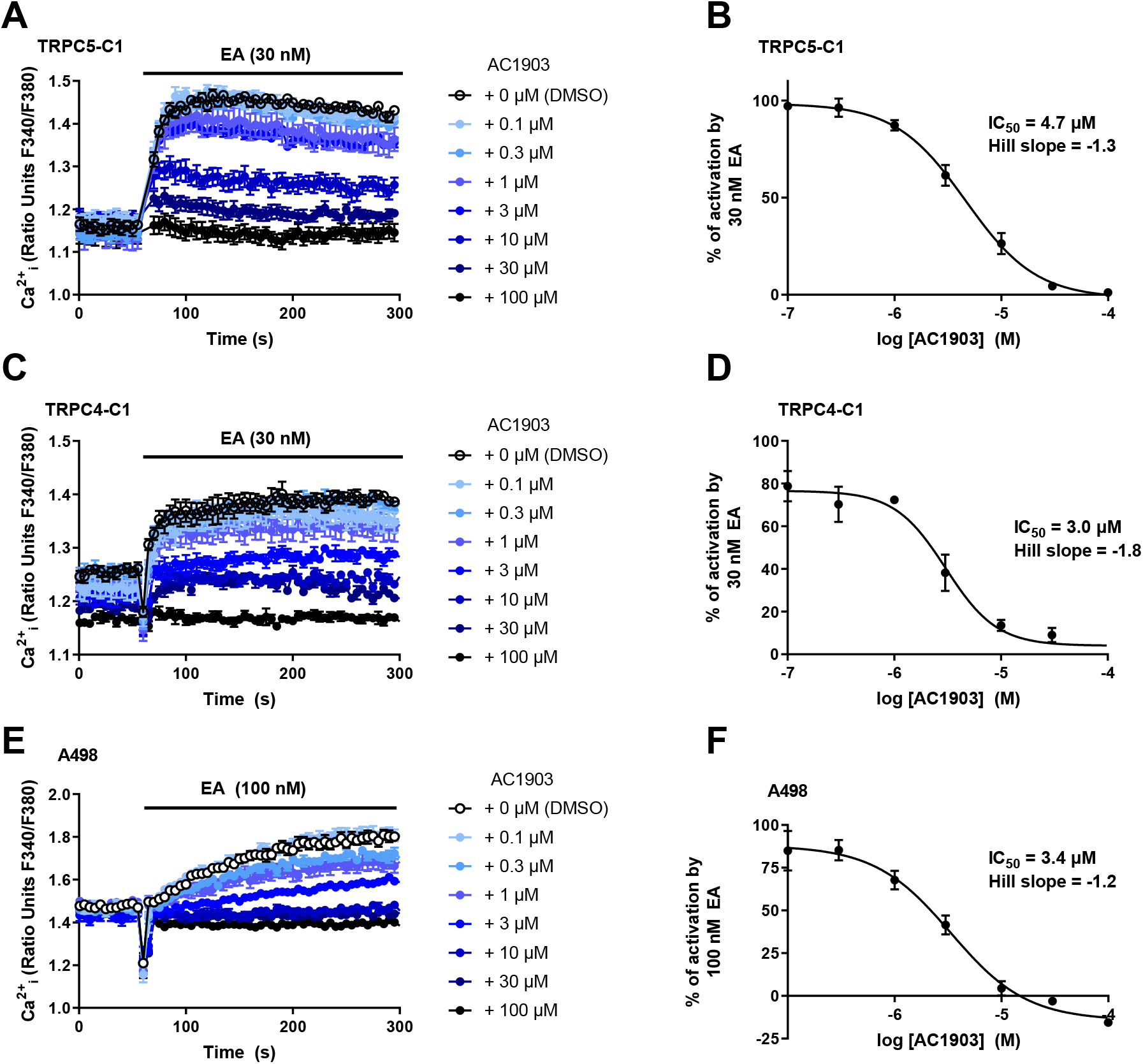
AC1903 inhibits heteromeric TRPC1/4/5 channels. A, C) Representative [Ca^2+^]_i_ measurements from a single 96-well plate (N = 6) showing inhibition of EA-mediated [Ca^2+^]_i_ responses by 0.1-100 μM AC1903 in (Tet+) HEK T-REx cells expressing concatemeric TRPC5-C1 (A) or concatemeric TRPC4-C1 (C). B,D) Concentration-response data for experiments in (A) and (C), showing mean responses ± SEM (n/N = 3-4/18-24). Responses were calculated at 250-295 s compared to [Ca^2+^]_i_ at baseline (0-55 s). E) Representative Ca^2+^ measurements from a single 96-well plate (N = 6) showing inhibition of EA-mediated [Ca^2+^]_i_ responses by 0.1-100 μM AC1903 in A498 cells. F) Concentration-response data for experiments in (E), showing mean responses ± SEM (n/N = 3/17-18). Responses were calculated at 250-300 s compared to [Ca^2+^]_i_ at baseline (0-55 s).

**Figure S10.**
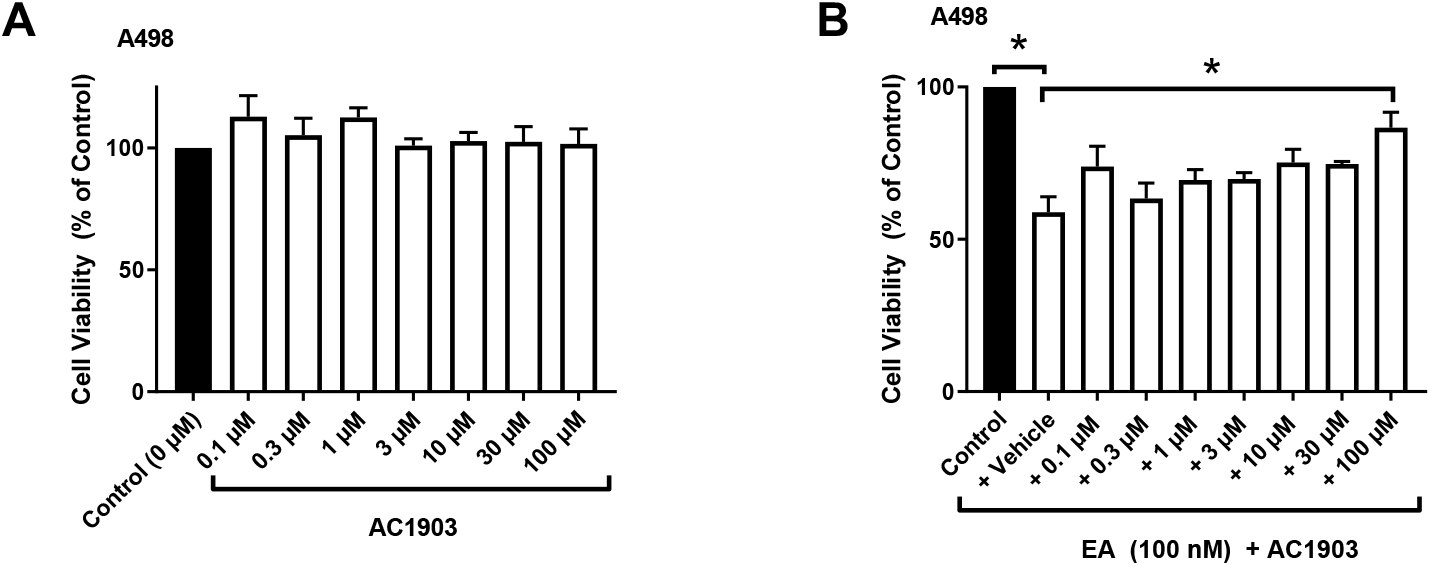
AC1903 is not toxic to A498 renal cancer cells but (at 100 μM) inhibits EA-mediated A498 cell death. A) Cell viability data for A498 cells treated with DMSO (control) or 0.1-100 μM AC1903 for 8 hours. Each condition was tested in triplicate. Cell viability was calculated as % of control, and the effect of AC1903 was compared to control (0 μM). Data are presented as mean ± SEM (n = 3). B) Cell viability data for A498 cells treated with DMSO (Control) or 100 nM EA in combination with either vehicle (DMSO) or 0.1-100 μM AC1903 for 8 hours. Each condition was tested in triplicate. Cell viability was calculated as % of control, and the effect of AC1903 was compared to EA + vehicle. Data are presented as mean ± SEM (n = 3).

**Figure S11.**
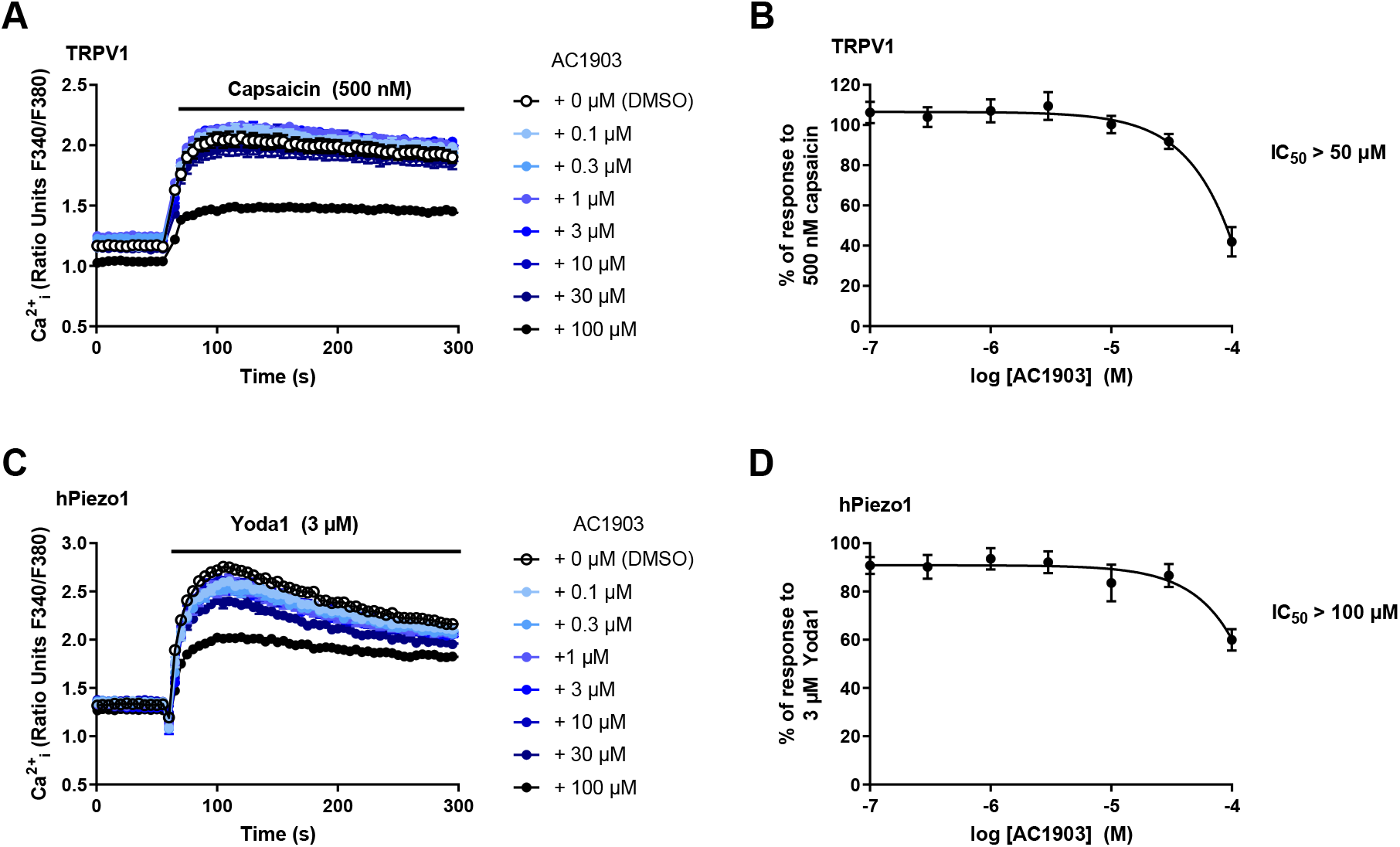
AC1903 (up to 30 μM) does not affect TRPV1 or hPiezo1 activation. A) Representative [Ca^2+^]_i_ responses from one 96-well plate (N = 6) showing the effect of DMSO or 0.1-100 μM AC1903 on capsaicin-mediated activation of TRPV1 expressed in HEK 293 cells. B) Concentration-response data for experiments in (A), showing mean responses ± SEM (n/N = 3/18). Responses were calculated at 95-100 s, compared to baseline at 0-55 s. C) Representative [Ca^2+^]_i_ responses from one 96-well plate (N = 6) showing the effect of DMSO or 0.1-100 μM AC1903 on Yoda1-mediated activation in HEK T-REx (Tet+) cells expressing hPiezo1. D) Concentration-response data for experiments in (C), showing mean responses ± SEM (n/N = 3/18). Responses were calculated at 95-115 s compared to [Ca^2+^]_i_ at baseline (0-55 s).

